# Characterizing nucleotide variation and expansion dynamics in human-specific variable number tandem repeats

**DOI:** 10.1101/2021.03.25.437092

**Authors:** Meredith M. Course, Arvis Sulovari, Kathryn Gudsnuk, Evan E. Eichler, Paul N. Valdmanis

**Affiliations:** Division of Medical Genetics, University of Washington School of Medicine, Seattle, WA, 98195, USA; Department of Genome Sciences, University of Washington, Seattle, WA, 98195, USA; Howard Hughes Medical Institute, University of Washington, Seattle, WA 98195, USA

**Keywords:** Variable number tandem repeats (VNTRs), long-read sequencing, human evolution, repeat instability, tandem repeat polymorphisms, repeat expansion, genome structure, single molecule, real-time (SMRT) sequencing

## Abstract

There are over 55,000 variable number tandem repeats (VNTRs) in the human genome, notable for both their striking polymorphism and mutability. Despite their role in human evolution and genomic variation, they have yet to be studied collectively and in detail, partially due to their large size, variability, and predominant location in non-coding regions. Here, we examine 467 VNTRs that are human-specific expansions, unique to one location in the genome, and not associated with retrotransposons. We leverage publicly available long-read genomes – including from the Human Genome Structural Variant Consortium – to ascertain the exact nucleotide composition of these VNTRs, and compare their composition of alleles. We then confirm repeat unit composition in over 3000 short-read samples from the 1000 Genomes Project. Our analysis reveals that these VNTRs contain remarkably structured repeat motif organization, modified by frequent deletion and duplication events. While overall VNTR compositions tend to remain similar between 1000 Genomes Project super-populations, we describe a notable exception with substantial differences in repeat composition (in *PCBP3*), as well as several VNTRs that are significantly different in length between super-populations (in *ART1, PROP1, WDR60*, and *LOC102723906*). We also observe that most of these VNTRs are expanded in archaic human genomes, yet remain stable in length between single generations. Collectively, our findings indicate that repeat motif variability, repeat composition, and repeat length are all informative modalities to consider when characterizing VNTRs and their contribution to genomic variation.

## Introduction

There are tens of thousands of variable number tandem repeats (VNTRs) in the human genome (Näslund et al. 2005), yet as a whole they remain uncharacterized. These VNTRs – that is, repeats with a repeat unit of seven base pairs (bp) or greater – are often too large or variable to be effectively captured using the short-read sequencing technologies typically used for whole genome sequencing. In addition, they are frequently located in non-coding or intergenic regions, which until recently have garnered less attention than genomic variants in coding regions. VNTRs, however, are highly mutable, suggesting that they play influential roles in evolutionary biology, and along with short tandem repeats (STRs; repeats with a repeat unit of fewer than seven bp), are a major source of human genetic diversity (Jeffreys et al. 1985; Berg et al. 2010; Hannan 2018). The recent advent of long-read sequencing has revealed that many VNTRs are much larger than the reference human genome suggests, and far more polymorphic. The few VNTRs that have been studied recently in more detail have provided insights into evolutionary history, replication mechanism, population structure, and disease. Characterizing more VNTRs at this higher resolution will continue to expand these insights.

Four VNTRs have recently been studied in detail, primarily due to their involvement in disease. A 25-bp VNTR in the intron of ATP binding cassette subfamily A member 7 (*ABCA7*) influences alternative splicing and is associated with Alzheimer’s disease (De Roeck et al. 2018). A 30-bp VNTR in calcium voltage-gated channel subunit alpha1 C (*CACNA1C*) exhibits varying repeat unit arrays correlated with “protective” or “risk” alleles in schizophrenia and bipolar disorder (Song et al. 2018). A 33-bp VNTR in the promoter of tribbles 3 homolog (*TRIB3*) is associated with *TRIB3* expression, and copy number of the repeat is correlated with certain disease-associated single nucleotide polymorphisms (Ord et al. 2020). We also identified a 69-bp repeat in WD repeat domain 7 (*WDR7*) associated with ALS (Course et al. 2020). A closer look at long-read sequenced genomes from geographically diverse samples indicated that this particular repeat expands via duplication events and a replication error called template switching. Furthermore, a small number of repeat units were unique to certain super-populations, and were also found in short-read datasets of ancient genomes. While examining this VNTR along with several others, we recognized that performing a similar analysis on a larger number of VNTRs could illuminate how these repeats vary and the mutational processes that have shaped them.

As these examples show, the VNTRs studied in detail have so far been studied one at a time and for a particular reason, like association with disease. Instead of continuing to study these repeats one-by-one, we decided to study a subset of them methodically and as a group. Doing so could answer questions about these VNTRs as a category of genomic variant, like their general characteristics as well as timing and patterns of expansion. Here, we look at a set of 467 VNTRs, chosen for the following characteristics: they exhibit human-specific expansion, they are not associated with retrotransposons, and they are unique to one location in the genome. These parameters select the VNTRs most likely to have expanded recently – and that may still be expanding – so we can observe their changes in different populations more readily. Furthermore, expansion of the same genomic segment in multiple places in the genome would be unlikely unless there were a retrotransposon driving it, so these parameters select for VNTRs that have expanded via other mechanisms. We then assess these or a subset of these VNTRs in ancestral human genomes as well as modern-day human genomes from the 1000 Genomes Project. We observe the similarities and differences of VNTRs in various super-populations, and their timescale of expansion. Finally, by taking a closer look at these genomes in long-read sequenced samples, we define several modalities of internal nucleotide pattering, which provides a useful framework for future VNTR analysis.

## Results

### Identifying VNTRs that are unique, non-retrotransposon associated, human-specific expansions

To generate a list of VNTRs of interest, we started with 1584 VNTRs that were recently described as having human-specific expansions (Sulovari et al. 2019). This list included both repeat expansions that arose *ab initio* (meaning they are only expanded in humans) and repeat expansions that are more expanded in humans than other primates. We then excluded VNTRs that were part of repetitive elements, like SINE-VNTR-Alu (SVA) repeats, and any repeats that were not unique to one location in the human genome (**Supp. Table 1**). These parameters allowed us to generate a list of 467 unique VNTRs with human-specific expansions that were unlikely to have arisen due to a retrotransposon. These VNTRs ranged in repeat unit size from 7-341 bp (mean = 40.1±28.6 bp; median = 34 bp; **Fig. 1A**). Average repeat copy number in the GRCh38 human reference genome ranged from 2-300.5 copies (mean = 39.8±40.4; median = 26.9). Comparing repeat motif size versus repeat copy number revealed an inverse correlation between motif size and copy number, fitting a log-log pattern of nonlinear regression (log-log slope = −0.74; **Fig. 1B**). Accordingly, average total repeat length in the GRCh38 human reference genome, which ranged from 65-14168 bp (mean = 1188±1078 bp; median = 945 bp) was not correlated with motif size (log-log slope = 0.16; **Fig. 1C**). This pattern was recapitulated in a separate dataset covering eight geographically diverse genomes obtained through PacBio small molecule, real-time (SMRT) long-read sequencing (Audano et al. 2019), in which the average total length of the repeats ranged from 86-8624 bp (mean = 1519±1099 bp, median = 1230 bp), and showed no correlation between repeat motif size and total length (log-log slope = 0.034; **Fig. 1D**). While these lengths were obtained through disparate sequencing methods, their similar means and standard deviations, as well as their similar nonlinear regression statistics, are concordant and indicate that both types of sequencing are useful for analyzing VNTR expansions.

**Figure 1.**
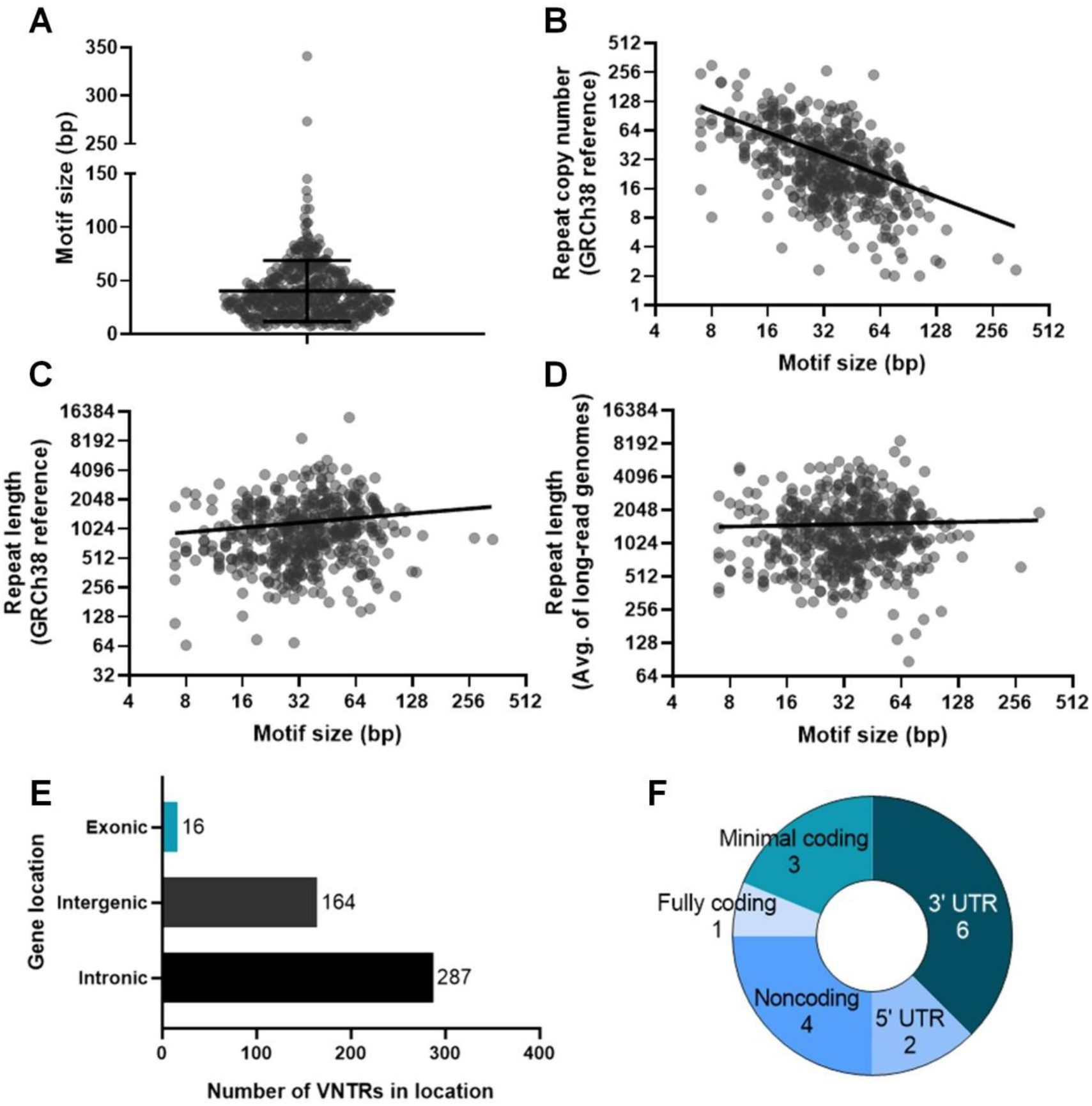
Characterization of 467 human-specific VNTR expansions assessed. (*A*) Violin plot of motif size for each VNTR. Lines show mean (40.1 bp) and standard deviation (±28.6 bp). (*B*) Scatter plot of motif size versus repeat copy number in the GRCh38 human reference genome. Axes are log_2_ and line is log-log (best-fit slope = −0.74). (*C*) Scatter plot of motif size versus total repeat length in the GRCh38 human reference genome. Axes are log_2_ and line is log-log (best-fit slope = 0.16). (*D*) Scatter plot of motif size versus average total repeat length in eight SMRT long-read sequenced genomes representing different super-populations. Axes are log2 and line is log-log (best-fit slope = 0.034). (*E*) Bar chart summarizing overall VNTR locations in the genome. (*F*) Pie chart breaking down specific locations of VNTRs in exons.

As for location in the genome, the vast majority of these VNTRs were in non-coding regions, with 287 intronic and 164 intergenic (**Fig. 1E**). Sixteen VNTRs were exonic, with four of these in noncoding exons, two in 5’ UTRs, and six in 3’ UTRs. Three overlapped coding regions by only 2, 5, or 15 bp out of a several hundred bp sequence (termed “minimal coding”), leaving only one VNTR that was fully contained within a coding region (**Fig. 1E, F**). This 60-bp VNTR is divisible by three bp, and resides in the gene *MUC1*, where it has previously been shown to play a role in medullary cystic kidney disease type 1 and other renal phenotypes (Kirby et al. 2013; Mukamel et al. 2021).

### Human-specific VNTR expansions are largely expanded in ancient genomes

We assessed the lengths of these 467 VNTRs in the short-read sequenced genomes of ancestral humans – specifically, an Altai Neanderthal genome (Prüfer et al. 2014) and a Denisovan genome (Meyer et al. 2012) (**Fig. 2A**). Of the 460 VNTRs that we successfully assessed, we found that only 16 VNTRs were not expanded in either genome, four were not expanded only in the Neanderthal genome, and three were not expanded only in the Denisovan genome (**Fig. 2B**). The number of expanded repeats may even be an under-estimate because of low coverage (especially of AT-rich sequences) and short read length in ancient genomes. Overall, we can conclude that the overwhelming majority of these 460 human-specific VNTR expansions are also expanded in ancient human genomes, suggesting that most expansions predate modern humans.

**Figure 2.**
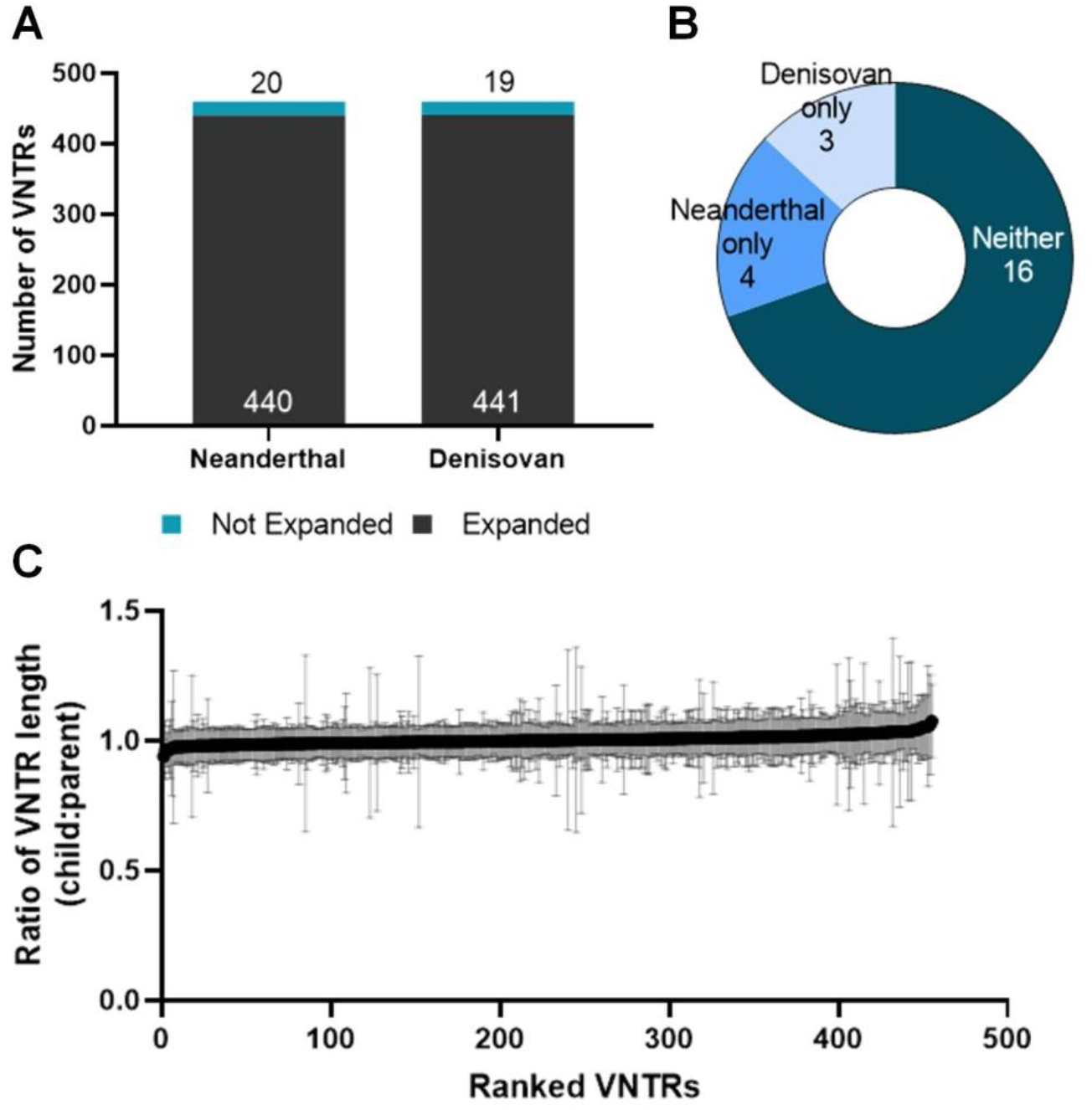
Timing of expansion in 467 human-specific VNTRs. (*A*) Bar chart showing number of VNTRs expanded (gray) or not expanded (blue) in Neanderthal and Denisovan genomes. 460 VNTRs were successfully assessed. (*B*) Pie chart breaking down VNTRs not expanded only in Neanderthal and/or Denisovan genomes, or neither. (*C*) XY plot showing mean (black dots) and standard deviation (gray lines) ratio of child VNTR length to average parent VNTR length. Data is from 585 trios from the 1000 Genomes Project. 455 VNTRs were successfully assessed, and are ranked by mean ratio on the x-axis.

### Human-specific VNTR expansions do not exhibit intergenerational expansion

To estimate the rate of VNTR expansion in modern-day humans, we observed these VNTRs in short-read sequenced trio datasets from the 1000 Genomes Project (Auton et al. 2015). Using 585 trios, we looked for events of expansion that occurred in one generation by comparing average repeat lengths in children versus parents. Ultimately, we observed that in virtually all of the 455 VNTRs successfully assessed, the average of the parents’ repeat lengths and the child’s repeat length remained the same (meaning the ratio between the two stayed at or near one). The lack of any obvious change in copy number between generations indicates that expansion in a single generation is rare for these VNTRs, and therefore they expand over a longer timescale (**Fig. 2C**).

### Internal sequence variation of human-specific VNTR expansions can be divided into three categories

To assess the internal nucleotide patterns in these VNTRs, we evaluated their repeat unit composition using existing long-read SMRT-sequenced datasets in a subset of 53 VNTRs. These VNTRs were chosen for having the greatest standard deviation in length in the original dataset (Sulovari et al. 2019), with the prediction that they would be the most likely to exhibit a variety of different alleles across the genomes assessed. We extracted genomic sequences of the 53 VNTRs from 15 individuals who had undergone whole genome long-read PacBio SMRT sequencing, representing the five super-populations from the 1000 Genomes Project (African, Admixed American, East Asian, European, and South Asian)(Audano et al. 2019). Reads were divided into individual repeat units based on their nucleotide composition, beginning with the most common repeat, extracted from the simple tandem repeats finder track (Benson 1999) from the UCSC Genome Browser (Kent et al. 2002). Each repeat was assigned a single letter code and then aligned. We then color-coded the varying repeat units to visualize their overall repeat pattern (**Fig. 3A**).

**Figure 3.**
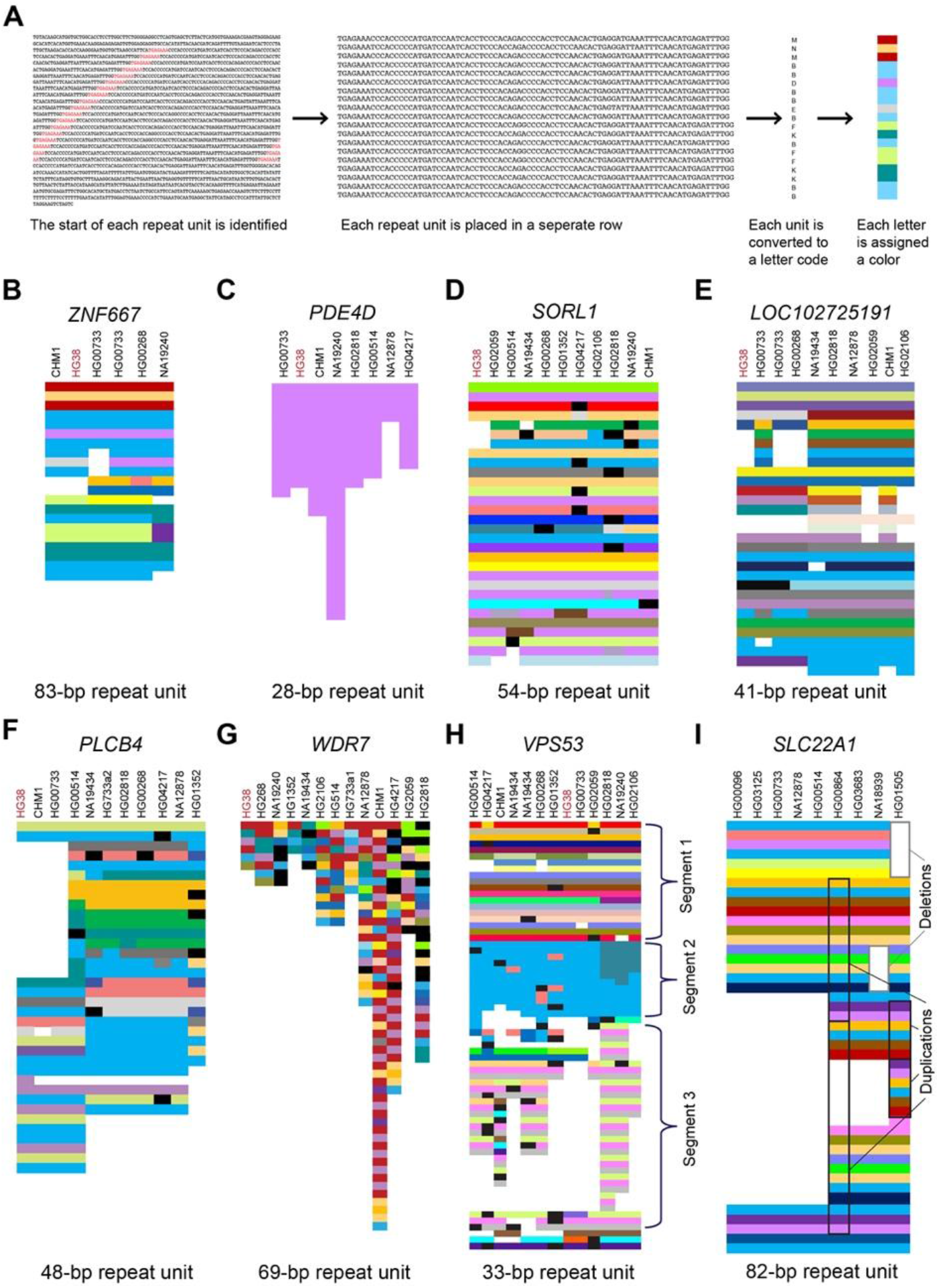
Composition plots illustrating modes of variability in VNTRs. (*A*) Schematic overview of how composition plots are generated from long-read sequencing. Example is from the CHM1 genome for the VNTR in *ZNF667. (B-H*) Composition plots showing varying patterns in example VNTRs, in (*B*) *ZNF667*, (*C*) *PDE4D*, (*D*) *SORL1*, (*E*) *LOC102725191*, (*F*) *PLCB4*, (*G*) *WDR7*, and (*H) VPS53.* At the top of each plot are listed the genomes from which the allele has been obtained, which were previously sequenced and published, and which represent geographically diverse populations (Audano et al. 2019). Black segments in the plots denote motifs that are private. (*I*) Composition plot for the VNTR in *SLC22A1*, with examples of duplication and deletion boxed in black and grey, respectively. The genomes used for this plot were previously sequenced and published (Ebert et al. 2021).

From this visualization, we identified that there are three chief modes of variability to consider when categorizing VNTR internal nucleotide patterns. Defining a motif as the sequence of one repeat unit and alleles as a series of motifs, we observed the variability in length between alleles, variability in sequence between motifs, and variability in motif organization within alleles. In the 53 VNTRs assessed, we observed that the majority contained several repeat motifs, and some variability in length between alleles, and little variability in sequence between alleles. A good representation of this common pattern is found in a VNTR with an 84-bp repeat in *ZNF667* (**Fig. 3B**). Some VNTRs showed far less variability in motifs, exemplified here by a 28-bp VNTR in *PDE4D* (**Fig. 3C**), which is “pure” or “uninterrupted,” though variable in allele length. Alternatively, in some VNTRs, most or even all repeat motifs were different from the one previous, such as in a 54-bp VNTR in *SORL1* (**Fig. 3D**); yet, despite its varied motifs, this and many similar VNTRs exhibit largely the same length and sequence between alleles. In another example, a 41-bp VNTR in *LOC102725191* (**Fig. 3E**), we observed the same pattern of variability in repeat motifs, but this time observed two clearly distinct alleles. Taking this pattern a step further, a 48-bp VNTR in *PLCB4* (**Fig. 3F**) exhibits about three distinct alleles, which each vary in length, as well.

We also observed a few rare VNTRs that were highly variable in all three modalities: motif, length, and allele sequence, like the previously-identified 69-bp repeat found in *WDR7* (**Fig. 3G**)(Course et al. 2020). For some VNTRs, we even observed some striking patterning of motifs within alleles. This scenario is best exemplified by a 33-bp repeat in *VPS53* (**Fig. 3H**) with a fixed length segment containing a variable internal sequence, followed by a variable length segment with fixed internal sequence, and finally a third variable length segment that repeats in groups of three motifs (**Fig. 3H**). Overall, we observed that most length and allelic differences were derived from duplications and deletions, as highlighted in an 82-bp repeat in *SLC22A1* (long-read genomes used to analyze *SLC22A1* were obtained from (Ebert et al. 2021))(**Fig. 3I**), rather than unit-by-unit changes. Detailed motif information for each of these repeats is available in **Supp. Table 2**.

### VNTR motifs in *PCBP3* are remarkably patterned across modern-day super-populations

We also looked at the distribution of the repeat units in the most variable VNTRs using the 1000 Genomes Project, to analyze population differences in repeat motif distribution in 25 geographically distinct modern-day populations (2504 individuals). For the most part, there were no notable differences in repeat motif distribution in these VNTRs. One exception, however, was a 66-bp repeat in intron 11 of Poly RC binding protein 3 (*PCBP3*), which was variable in all three parameters: motifs, allele length, and allele sequence composition. A total of eight of the 66-bp in the repeat unit were variable, and these variable positions result in 38 total primary repeat motifs (**Fig. 4A**). A number of these repeat motifs were also observed in one Denisovan (Meyer et al. 2012) and three Neanderthal genomes (Prüfer et al. 2014, 2017; Mafessoni et al. 2020). The repeat motif sequence varied between alleles, although what similarities existed partially clustered by super-population (**Fig. 4B**). The frequency of these repeat motifs differed across super-populations, with a particularly noticeable inverse relationship between repeat motifs abundant in the African super-population versus the East Asian super-population (**Fig. 4C**).

**Figure 4.**
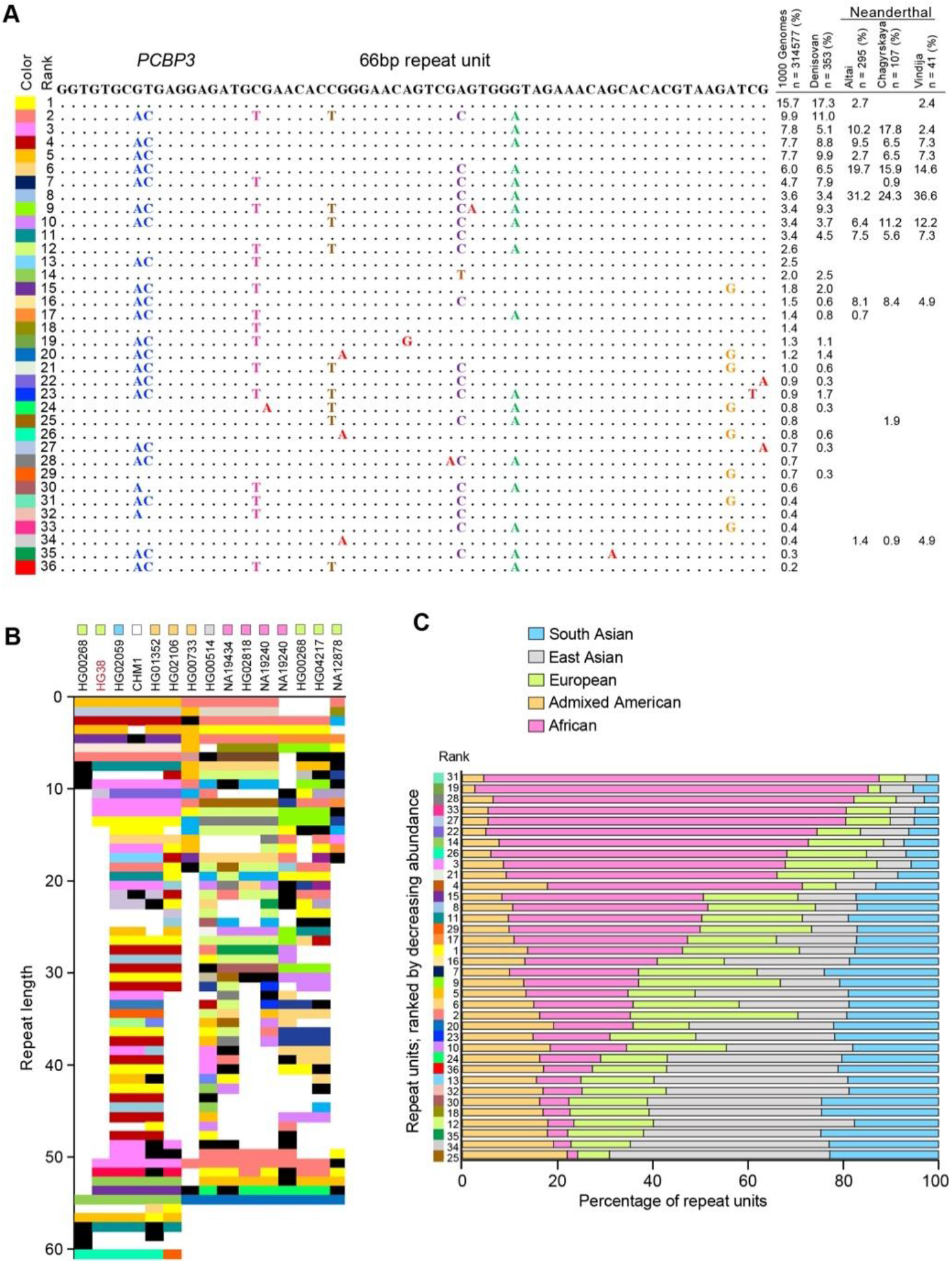
A VNTR in *PCBP3* shows repeat motif differences in modern-day super-populations. (*A*) *PCBP3* repeat motifs with variable positions highlighted. At left is the assigned color-code for each motif. At right is the relative abundance of each motif in the 1000 Genomes Project and in ancient genomes. (*B*) Composition plot of the *PCBP3* VNTR in geographically diverse populations. At the top of the plot is listed the genomes from which the allele has been obtained, which were previously sequenced and published (Audano et al. 2019). Black segments in the plot denote motifs that are private. (*C*) Cumulative frequency of repeat motifs in *PCBP3* across super-populations. Repeat motifs are ordered by decreasing abundance in the African super-population, and numbers on the y-axis correspond to their global ranked abundance.

### Some human-specific VNTR expansion lengths are significantly different across modern-day super-populations

We then used the same 1000 Genomes Project dataset to analyze the differences in length of all 467 VNTRs. We calculated the length of each repeat in these samples, merging combined reads for each of the five super-populations. After estimating the average length of the VNTRs across these populations, we generated volcano plots to determine which VNTRs had significant length differences between super-populations (**Fig. 5**). We observed that some super-populations had more similar average VNTR lengths than others. For example, European and Admixed American populations showed strong concordance between repeat length, and South Asian and East Asian populations had only one VNTR that was a notable outlier. From these volcano plots, we chose the top four differentially expanded VNTRs (by both p-value and log_2_-fold change) to observe further, in: *LOC102723906*, PROP Paired-Like Homeobox 1 (*PROP1*), ADP-ribosyltransferase 1 (*ART1*), and WD-repeat containing protein 60 (*WDR60*).

**Figure 5.**
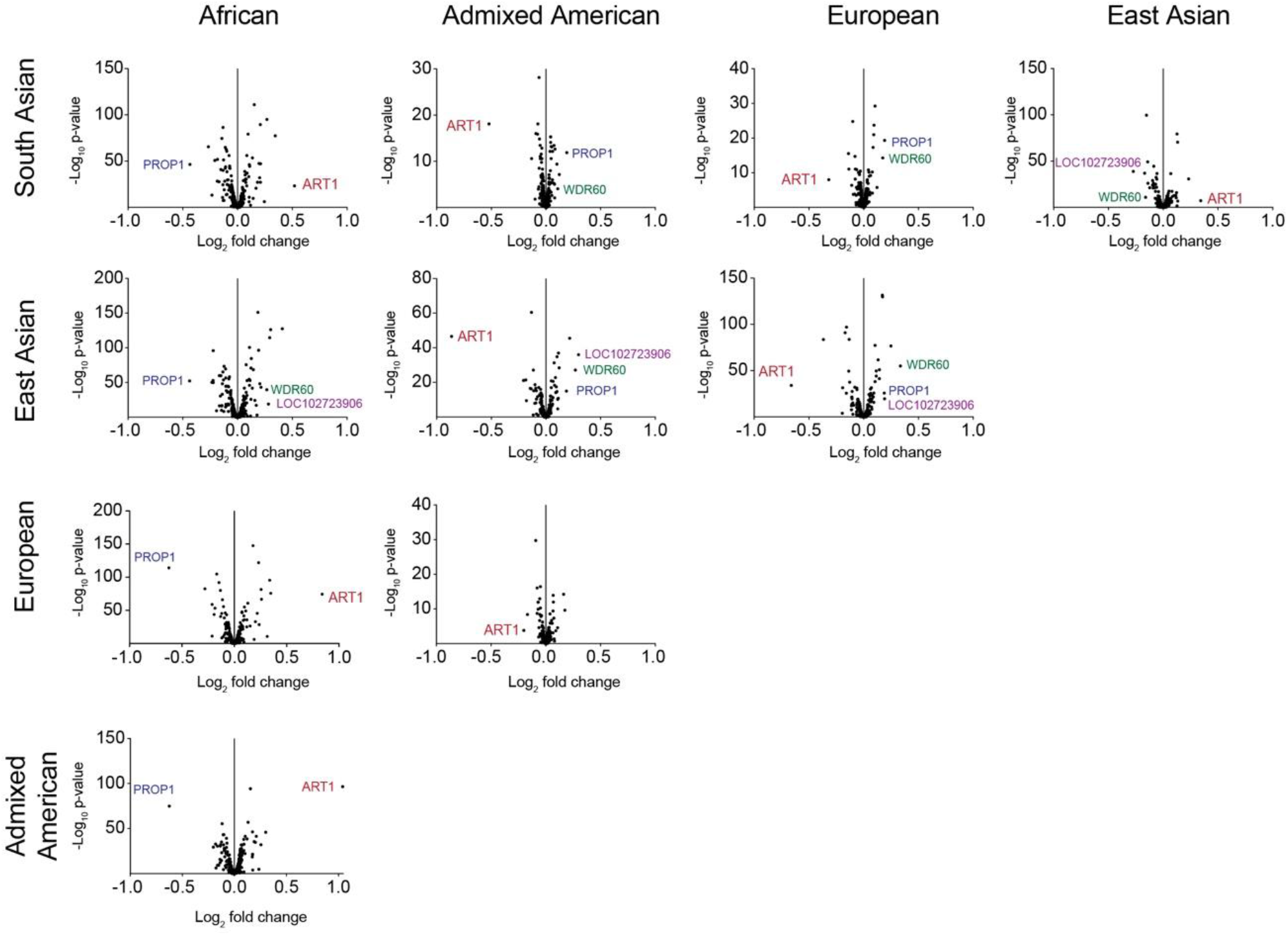
Comparing VNTR lengths across modern-day super-populations. Volcano plots showing pair-wise comparisons of average VNTR lengths between super-populations from the 1000 Genomes Project. The VNTRs with the greatest length differences (as determined by DESeq2) are labeled by the nearest gene or gene in which they reside, and were determined based on both log2-fold change and p-value. Trial size for each super-population is 347-660 individuals.

We compared both the pattern of individual data points as well as the cumulative abundance of the repeat lengths in all super-populations (**Fig. 6A-D**). One-way ANOVAs to compare each population for each VNTR all gave p < 0.0001, and subsequent Tukey’s multiple comparison tests corroborated the significant differences we observed in **Fig. 6** using DESeq2. Comparing these distributions, we saw that the VNTR in *LOC102723906* is longer in East Asian than European and African populations, though samples of African ancestry show a larger range of allele lengths (**Fig. 6A**). The cumulative plot mirrors the right-tailed distribution observed in African samples, and reveals an unusual “trimodal distribution” in the remaining super-populations (**Fig. 6A**). The VNTR in *PROP1* shows longer repeats in the African super-population than in any other (**Fig. 6B**), while the VNTR in *ART1* shows the opposite: fewer long repeats appearing in the Africa super-population, as compared to the other populations (**Fig. 6C**). The *ART1* cumulative plot shows an overall right-tailed distribution of repeat copy number. In addition, the VNTR in *ART1* shows a distinct gap in values in some of the super-populations, akin to a bimodal distribution (**Fig. 6C**). Finally, the VNTR in *WDR60* shows a more normal distribution across the super-populations, with longer repeats in the East Asian super-population (**Fig. 6D**).

**Figure 6.**
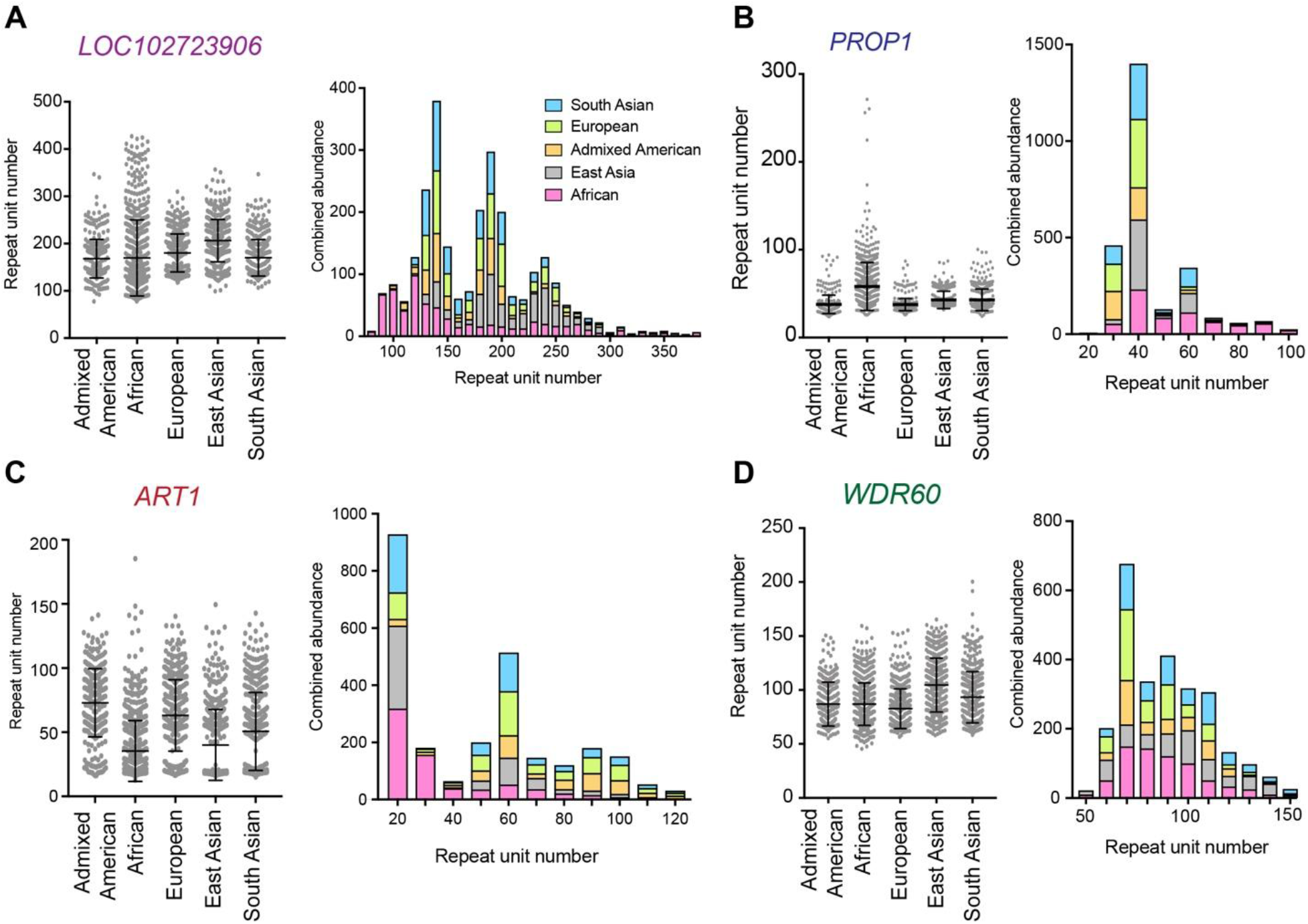
Length differences in top four differentially expressed VNTRs in modern-day super-populations. (*A-D*) Individual VNTR copy numbers plus mean and standard deviation for each superpopulation (left) and cumulative abundance binned into groups of 10 repeat units (right) for VNTRs in (*A*) *LOC102723906*, (*B*) *PROP1, (C) ART1*, and (*D*) *WDR60.* Trial size for each super-population is 347-660. One-Way ANOVAs gave p < 0.0001 for each comparison of super-populations for each VNTR.

### Differentially-expressed human-specific VNTRs expanded at different time points

We also determined the evolutionary timing of these four differentially-expressed expansions by observing them in the reference genomes of non-human primates. We found that they did not all initially expand around the same time. Instead, the VNTR in *WDR60* expanded between the branch point of New World monkeys and gibbons (the macaque genome has a portion of the sequence present), the VNTR in *LOC102723906* expanded between gibbons and orangutans, and the VNTRs in *PROP1* and *ART1* expanded between the branching of chimpanzees and hominins. All of these VNTRs are expanded in the Denisovan and Neanderthal genomes.

### Composition of differentially-expressed human-specific VNTRs explains findings in short-read sequencing

We then visualized the pattern of repeat motifs for these VNTRs as described in **Fig. 3A**, this time using the 32 available PacBio HiFi SMRT-sequenced genomes available through the Human Genome Structural Variant Consortium (HGSVC)(Ebert et al. 2021), ultimately providing up to 64 alleles. This analysis revealed that the unusual trimodal distribution observed in the cumulative plot for the VNTR in *LOC102723906* is explained by the three predominant alleles of three differing sizes (averaging 58, 109, and 253 repeat units) found by SMRT sequencing (**Fig. 7A**). Long-read sequencing also revealed a highly unusual internal structure in *PROP1*, in which each repeat motif itself contains a TG-stretch that can vary widely in length – between 7 and 26 repeat copies – like an STR within a VNTR (**Fig. 7B**). This TG-stretch is likely the main cause of its length variability. Long-read sequencing also explained the distinct gap in values observed in *ART1*, showing that there are three predominant alleles, two of which are similarly short (25 and 41 repeat units), and one of which is quite long (381 repeat units)(**Fig. 7C**). Larger expansions appear to occur as duplication events in blocks of 45-50 repeat units. Finally, the even distribution of lengths in *WDR60* is echoed by a varied distribution of alleles in long-read sequencing (**Fig. 7D**), not to mention over 100 different motifs. Overall, all of these four VNTRs have markedly variable internal nucleotide sequence, a couple of major alleles, and except for the VNTR in *PROP1*, their major length variation is due to clear deletion or duplication events. It is worth noting that these VNTRs tend to be longer and more variable than other VNTRs in this dataset, which may be related to the fact that they are also the most different in length between super-populations.

**Figure 7.**
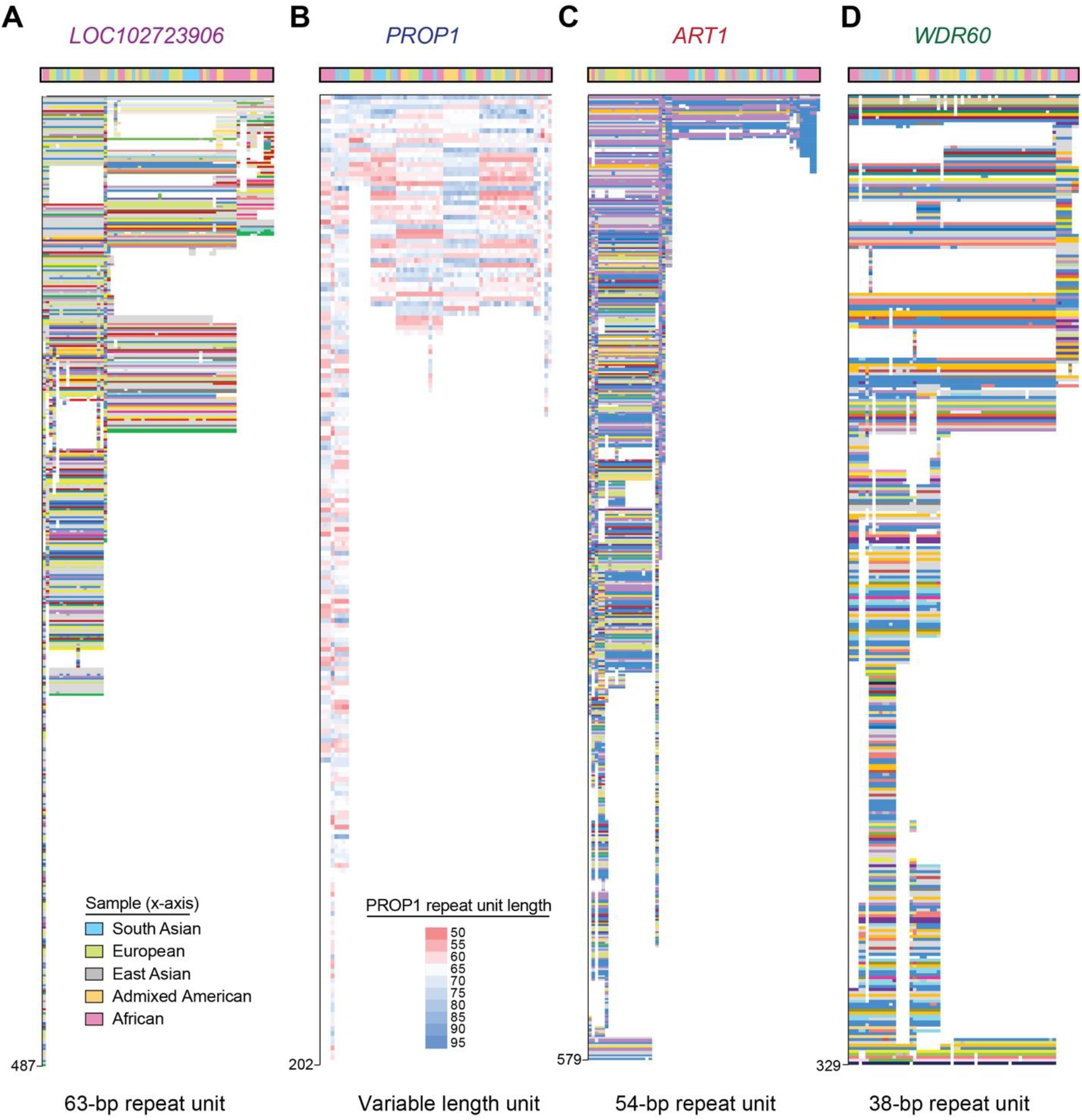
Composition plots of top four differentially expressed VNTRs in modern-day super-populations. Composition plots for VNTRs in (*A*) *LOC102723906*, (*B*) *PROP1*, (*C*) *ART1*, and (*D*) *WDR60.* The colors at the top of the plot denote the super-population from which the alleles were obtained (see key), which were previously sequenced and published (Ebert et al. 2021). Grey segments in the plot denote motifs that are rare or private. Y-axis shows length of the repeat in bp. The heat map legend in (*B*) denotes the length of each repeat found in the *PROP1* VNTR, which has been plotted based on this unique feature, instead of the motif structure used for the other VNTRs.

## Discussion

In this study, we observed the patterns, timing of expansion, and geographical distribution of 467 VNTRs that are unique, non-retrotransposon associated, and expanded specifically in humans. These particular characteristics were chosen to enrich for VNTRs that were likely to have expanded recently and via mechanisms other than association with repetitive elements. Even by generating this limited subset, we still observed a wide variability in repeat motif size and sequence, repeat length, internal sequence variability, and allele variability (**Fig. 1, 3**).

In considering the timing of expansion of these VNTRs, we first found that the overwhelming majority are also expanded in ancient human genomes, therefore most expansions predate modern humans. (**Fig. 2A, B**). Human-specific expansion refers to both rare repeat expansions that arose *ab initio* (meaning they are only expanded in humans) as well as repeat expansions that are more expanded in humans than other primates. When we looked closer at the four VNTRs most different in length across super-populations, we saw that they initially appeared at three different points in the primate phylogenetic tree, so there is no generalized pattern or moment that explains this initial expansion as a group. Finally, looking at modern-day humans, we found that none of the trios in the 1000 Genomes Project show obvious expansion over a single generation timescale. This intergenerational stability suggests that the VNTRs expand over a longer timescale, and therefore the events that lead to their expansion occur less frequently (**Fig. 2C**). This observation is consistent with what we previously observed using SMRT long-read sequencing in a very large pedigree for the *WDR7* VNTR (Course et al. 2020). It is also consistent with the negative relationship we observed between motif size and copy number (**Fig. 1B**), which indicates that the longer a motif length is, the less likely it is to expand rapidly. That said, observing VNTRs in pedigrees has long been suggested as a way to differentiate between allelic variability due to homologous recombination versus unequal sister chromatid exchange, and looking for rare mutation events is still worthwhile (Jeffreys et al. 1985).

As for the geographical distribution of these VNTRs in modern-day humans, we see that some populations share more similar overall VNTR lengths than others (i.e. European and Admixed American super-populations share few VNTRs with a large log2-fold change), while there are many repeats whose lengths are significantly different between populations (**Fig. 5**). The VNTRs with the greatest length differences were in *ART1, PROP1, WDR60*, and *LOC102723906.* All of these VNTRs were notably variable in internal nucleotide composition, but in most other ways they differed from one another, including in their length distribution patterns (**Fig. 6**) and their allele patterns revealed by long-read sequencing (**Fig. 7**). Long-read sequencing analysis of these VNTRs explained the patterns we observed from short-read data, with the primary alleles accounting for the trimodal distribution pattern observed for the VNTR in *LOC102723906* (**Fig. 6A**), and the gap in lengths observed for the VNTR in *ART1* (**Fig. 6C**), for example. The reason why some populations have a higher frequency of certain expanded alleles is unclear, but it is consistent with population bottlenecks followed by drift of alleles that are possibly non- or weakly deleterious. This analysis also indicates that many VNTRs continue to expand in certain super-populations. While length may change noticeably between some super-populations, the repeat motifs themselves generally do not, perhaps because the variation originated a long time ago. One intriguing exception to this observation is a VNTR in *PCBP3*, which exhibits an almost inverse correlation between repeat motifs common in the African super-population and those in the East Asian super-population (**Fig. 4C**).

We used existing SMRT long-read datasets to resolve the internal nucleotide patterns of 53 of these VNTRs, chosen for their high standard deviation in length (Sulovari et al. 2019), in addition to the four observed in the context of population structure. In doing so, we have determined that the primary considerations for VNTR analysis are: variability in motif organization within alleles, variability in motif sequence between alleles, and variability in length between alleles. Most VNTR analysis has revolved around assessing variability in length between alleles; however, we hope to illustrate here that the remarkable sequence compositions of VNTRs are a critical consideration, as well. Recent examples already indicate that sequence composition of VNTRs can influence disease state (Song et al. 2018; De Roeck et al. 2018), in addition to revealing population differences and expansion patterns (Course et al. 2020). Even in this subset of VNTRs, we observe a wide range of patterns. Some repeats are quite variable in allele length, while others seem to have only a few alleles fixed at certain lengths. Some repeats are not variable in sequence, while some are partially variable, and some others still are extremely variable, based on the number of different repeat units observed (**Fig. 3**). VNTRs can even have sections that represent more than one of these categories (ex. **Fig. 3H**). We did observe that most of the VNTRs lacked high allele variance across super-populations, which makes sense in light of the fact that most are already expanded in ancient genomes.

The primary force driving differences between alleles – both sequence and length – appears to be deletions and duplications (ex. **Fig. 3I**), which could occur anywhere in the repeat, and are possibly the result of homologous recombination or unequal sister chromatid exchange. Compared to the VNTR in *WDR7* that we had previously studied (Course et al. 2020)(**Fig. 3G**), none the VNTRs whose internal nucleotide patterns that we assessed showed as clear a directionality of expansion. This observation corroborates what has been seen for the VNTRs in *ABCA7* (De Roeck et al. 2018), which is dynamically expanded in non-human primates and has much more internal variability, and in *CACNA1A* (Song et al. 2018), which has variability primarily in two regions in the middle of the repeat. Some of the earliest studies of VNTR evolution pointed towards both mitotic/meiotic recombination and replication slippage as influencing VNTR expansion and variability (Jeffreys et al. 1985). Repeat directionality (or polarity) has been observed in STRs, where the directionality played a role in instability of the repeat (Eichler et al. 1995). As more genomes undergo long-read sequencing, we may find additional examples of directionality in these longer VNTRs.

After observing the wide range of motif size in this set of VNTRs (7-341 bp), encompassing a mean size of 40 bp, together with the wide range of internal sequence variability and patterning, we submit that the current definition of a VNTR as a tandem repeat with seven bp motif or larger could benefit from more comprehensive categorization. We initially chose this set of VNTRs by selecting for parameters like human-specific expansion, but it may be more useful going forward to analyze VNTRs together in groups of physical characteristics, like motif size, motif variability, and allele variability. This kind of grouping is more likely to reveal common mechanisms of expansion and evolution.

Overall, these human-specific VNTR expansions are generally found in non-coding regions, are already expanded in ancient human genomes, and remain stable between single generations. They tend to have some variability in their motifs, but less variability between alleles, with their primary source of allelic variation driven by deletions and duplications. That said, there are exceptions to each of these generalizations, and what is clearly different between each VNTR is their motif size, which ranges widely and correlates inversely with copy number; their patterns of motif variability and sequence composition; and when they first arise in non-human primates. Sub-categorizing VNTRs with these considerations in mind, and observing them in more detail with long-read sequencing, may lead to uncovering larger patterns that explain their expansion dynamics. The number of publicly available long-read genomes is expected to increase significantly in the next several years, and this study provides a framework for conducting further analysis.

## Methods

### VNTR selection

We started with a list of 1584 tandem repeats categorized by their expansion specifically in humans (Sulovari et al. 2019). From this list, we excluded SVA retrotransposons, and any repeats with a motif size less than 7 nt. These selection criteria yielded 467 VNTRs with unique matches in the human genome.

### VNTR gene location

To determine the locations of the VNTRs, we started with previously published data (Sulovari et al. 2019). We then manually inspected VNTRs that were within or close to exons and compared VNTR location to GENCODE v32 transcripts in the UCSC Genome Browser (Kent et al. 2002).

### Repeat length estimation

We adapted previous methods (Course et al. 2020; Song et al. 2018) to estimate read depth using short-read whole genome sequencing data. Reads were counted that mapped to the repeat, compared to three sets of 100 kb windows of genomic sequence. The fraction of enrichment or depletion of reads was used to calculate the estimated length of each VNTR compared to the reference human genome (GRCh38).

Raw data for ancient DNA calculations were obtained from published genomes of Altai Neanderthal (Prüfer et al. 2014) and Denisovan (Meyer et al. 2012) samples. We converted the coordinates of each VNTR from GRCh38 to hg19 using the liftOver tool in the UCSC Genome Browser, then queried the ancient genomes for the number of matches within those coordinates padded by 2kb on either side of the VNTR repeat sequence as listed by Tandem Repeat Finder (Benson 1999). Of the 467 VNTRs, seven did not have corresponding matches in hg19. Regions corresponding to each VNTR were converted to SAM files and individual reads were queried for the presence of a complete repeat unit. For *PCBP3*, we also searched for repeat unit sequence matches from DNA reads for Chagyrskaya and Vindija Neanderthal Genomes (Prüfer et al. 2017; Mafessoni et al. 2020). Total counts for each repeat unit were aggregated across each super-population, and used to identify their relative abundance.

Raw data for modern-day DNA calculations were obtained from the 1000 Genomes Project (Auton et al. 2015). Reads that mapped to each VNTR were extracted from BAM alignment files for each individual. The number of reads that corresponded to each repeat was counted across each sample and normalized first to the length of the repeat in the GRCh38 human genome and then to the read density across three separate 100kb segments of DNA that did not contain a VNTR (which were confirmed to have consistent read density estimates across all three bins). This number of aligned reads from each sample was used as input for the DESeq2 program (Love et al. 2014) to identify differentially expanded repeats between super-populations.

### Long-read sequencing analysis

We obtained 15 PacBio SMRT-sequenced genomes from (Audano et al. 2019), and 32 from the HGSVC (Ebert et al. 2021). We then aligned and visualized the VNTR alleles from these datasets as described previously (**Fig. 3A**)(Course et al. 2020). Briefly, after extracting the repeat sequence along with flanking intronic sequence, each repeat unit per VNTR was assigned a single letter or number code. Each letter was then converted to its own unique color to improve visualization when aligning the series of repeat units for each sample. We then organized the repeat sequences based on their similarity, including creating gaps to shift units down to match the arrangement of neighboring alleles. Perhaps because of their repetitive nature, we did not always identify each repeat in every sequenced individual. Samples were not included if their repeat abutted the end of a sequence contig, which was a relatively common occurrence.

### Phylogenetic analysis

Presence or absence of the VNTR repeat unit of interest was observed by extracting sequences in non-human primates in the UCSC Genome Browser. Using the “View in other genomes (convert)” function in the UCSC genome, we extracted sequences of 10 non-human primates to determine the presence of one or more copies of the repeat motif.

### Statistical analyses

Statistical analyses were performed using Prism 8.0.1 (GraphPad Software). Log-log line slopes were determined by least squares regression. Volcano plots were generated by graphing log2 fold change against negative log_10_ p-value on an XY plot. A one-way analysis of variance (ANOVA) test was used to compare groups greater than two, followed by Tukey’s multiple comparisons test when the ANOVA gave p < 0.05.

## Data Access

Whole genome long-read sequence data used in this study was obtained from the NCBI BioProject database (https://www.ncbi.nlm.nih.gov/bioproject) with accessions NCBI:PRJNA246220 (CHM1), NCBI:PRJNA300843 (HG00514), NCBI:PRJNA300840 (HG00733), NCBI:PRJNA288807 (NA19240), NCBI:PRJNA339722 (HG02818), NCBI:PRJNA385272 (NA19434), NCBI:PRJNA339719 (HG01352), NCBI:PRJNA339726 (HG02059), NCBI:PRJNA323611 (NA12878), NCBI:PRJNA481794 (HG04217), NCBI:PRJNA480858 (HG02106), and NCBI:PRJNA480712 (HG00268). Data from the HGSVC is available from https://www.internationalgenome.org/data-portal/data-collection/hgsvc2. 1000 Genomes project samples and short-read sequence data is available from https://www.internationalgenome.org/data#download. Data for Altai Neanderthal and Denisovan genomes can be accessed from ENA archives ERP002097 and ERP001519, respectively. Samples for Vindija and Chagyrskaya Neanderthals were accessed from http://cdna.eva.mpg.de/neandertal/Vindija/ and http://cdna.eva.mpg.de/neandertal/Chagyrskaya/, respectively.

## Acknowledgments

This work was supported by the National Institute of General Medical Sciences (5T32GM007454-38 to M.M.C.), and the National Human Genome Research Institute (HG010169 to E.E.E.). E.E.E. is an investigator of the Howard Hughes Medical Institute. P.N.V. is supported in part by the Robert F. Schoeni Award for Research from the Ann Arbor Active Against ALS. We acknowledge the support of all members of the Valdmanis lab. Finally, we thank all individuals who donated biospecimens for their willingness to contribute to scientific research.

## Competing Interest Statement

The authors declare that they have no competing interests.

## Notes

### Competing Interest Statement

The authors have declared no competing interest.

### Summary of Updates

This version of the manuscript includes Supplementary Tables 1 and 2

